# Identification of FMRP target genes expressed in corticogenesis: implication for common phenotypes among neurodevelopmental disorders

**DOI:** 10.1101/769026

**Authors:** Cristine R. Casingal, Takako Kikkawa, Hitoshi Inada, Noriko Osumi

## Abstract

Fragile X mental retardation protein (FMRP) is encoded by *FMR1* gene that is responsible for Fragile X Syndrome (FXS) showing intellectual disability and autism spectrum disorder. FMRP is an RNA binding protein highly expressed in the brain. Although several target genes for FMRP have been identified, limited studies have suggested the role of FMRP in corticogenesis. Here we performed RNA immunoprecipitation sequencing against the murine embryonic neocortex, and identified 124 genes as potential FMRP mRNA targets. We found 48 of these genes as overlapped with autism-related genes, which were categorized in four functional groups: “transcriptional regulation”, “regulation of actin cytoskeleton”, “ubiquitin-mediated proteolysis” and “calcium signaling pathway”. Four of these genes showed significant difference in expression in the cortical primordium of *Fmr1-*KO mice; *Huwe1* and *Kat6a* increased, while *Kmt2c* and *Apc* decreased. Although the change in expression of these four genes was relatively small, these subtle changes due to dysregulated transcription could collectively contribute to impaired corticogenesis to cause phenotypes of FXS. Investigating the transcriptional control of FMRP on its mRNA targets may provide new insight to understand neurodevelopmental pathogenesis of FXS.

## INTRODUCTION

Fragile X Syndrome (FXS) is an X-linked neurodevelopmental disorder that causes intellectual disability as well as behavioral deficits. Among genetically caused cases of autism spectrum disorder (ASD), the most frequent neurodevelopmental disorder, 1-2% of the patients exhibit FXS (Abrahams and Geschwind, 2008; Garber *et al.*, 2008; Dahlhaus, 2018). Patients with FXS have an abnormal expansion of CGG repeats in the 5’-untranslated region of the gene, *fragile X mental retardation* (*FMR1*). This results in the hypermethylation and transcriptional silencing of the gene, which lead to the loss of its product fragile X mental retardation 1 protein (FMRP) (Verkerk *et al*., 1991; Ashley Jr. *et al*., 1993; Bakker *et al*., 1994; Crawford *et al*., 2001; Garber *et al*., 2008). FMRP is a polyribosome-associated RNA-binding protein (RBP) that selectively targets specific mRNAs, regulates its translation, transport, and stability, as well as histone modification and chromatin remodeling (Bassell and Warren, 2008; De Rubeis and Bagni, 2010; Alpatov *et al.*, 2014; Korb *et al.*, 2017). Hence, FMRP is a multifunctional protein that could be involved in diverse processes, not limited to the mRNA lifecycle.

FMRP is widely expressed in the embryonic and adult brain. During corticogenesis, it is expressed in neural stem cells (or radial glial cells, RGCs), localizing at the apical and basal endfeet of the RGCs (Saffary and Xie, 2011; La Fata *et al.*, 2014; Pilaz *et al.*, 2016), regulates transition from RGC to intermediate progenitor (Saffary and Xie, 2011), and affects neuronal migration and cortical circuitry (La Fata *et al.*, 2014). Postnatally, FMRP is localized in the cell body, proximal dendrites and axons of neurons (Darnell *et al.*, 2011; Castrén, 2016), and plays profound regulatory roles in the synaptic function and neuronal plasticity (Bassell and Warren, 2008; De Rubeis and Bagni, 2010) through interaction with transcripts that encode pre- and postsynaptic proteins, and also regulation of mRNA trafficking into dendrites (Abrahams and Geschwind, 2008; Bassell and Warren, 2008; Abrahams *et al.*, 2013). Altogether, FMRP has multifunctional roles at distinct times in brain development.

Since the discovery of FMRP, a large effort has been made to characterize targets of FMRP using several methods including RNA immunoprecipitation (RIP) followed by microarray analysis (Brown *et al.*, 2001; Pilaz *et al.*, 2016), crosslinking immunoprecipitation followed by high-throughput sequencing (Darnell *et al.*, 2011), and photoactivetable ribboneoside-enhanced crosslinking and immunoprecipitation (Ascano *et al.*, 2012). However, an inclusive research to identify FMRP mRNA targets and their roles is limited during brain development compared with those in postnatal stages.

In this study, we identified a large number of FMRP mRNA targets using RIP high-throughput sequencing in the mouse embryonic cortex. These FMRP target genes showed highly significant overlap with ASD-related genes. We found that these molecules were associated with “transcriptional regulation”, “regulation of actin cytoskeleton”, “ubiquitin-mediated proteolysis” and “calcium signaling pathway”. Among the candidate FMRP mRNA targets, the expression levels of *Huwe1* and *Kat6a* were increased and those of *Kmt2c* and *Apc* were decreased in the *Fmr1*-knockout (KO) mice (Bakker *et al.*, 1994), implying a possibility that FMRP may regulate its mRNA targets at the level of transcription. The dysfunction of FMRP may thus lead to changes the developmental program during corticogenesis in multiple ways.

## RESULTS

### FMRP expression in the mouse embryonic cortex

We first tried to confirm the expression of FMRP in the neocortical primordium of wild type (WT) mice at embryonic day (E) 14.5 when massive neurogenesis occurs. FMRP was accumulated at apical and pial (basal) surface areas of the cortical primordia (Figure 1A-C). To confirm the detailed expression of FMRP in the surface areas of the cortex, we labeled the RGCs using *in utero* electroporation with pCAG-EGFP at E13.5 mice, and sacrificed at E14.5. Immunostaining revealed that the FMRP protein was overlapped with GFP in the apical and basal endfeet of the RGCs (Figure 1D-I). We also observed strong expression of FMRP in the cortical plate consisting with immature neurons (Figure 1A-C), which has not been reported in the above-mentioned previous studies (Saffary and Xie, 2011; La Fata *et al.*, 2014; Pilaz *et al.*, 2016). This may be attributed to difference in antibodies used, although it is not surprising that FMRP is indeed expressed in neurons of adult mice (Darnell *et al.*, 2011; Maurin *et al.*, 2018).

**Figure 1.**
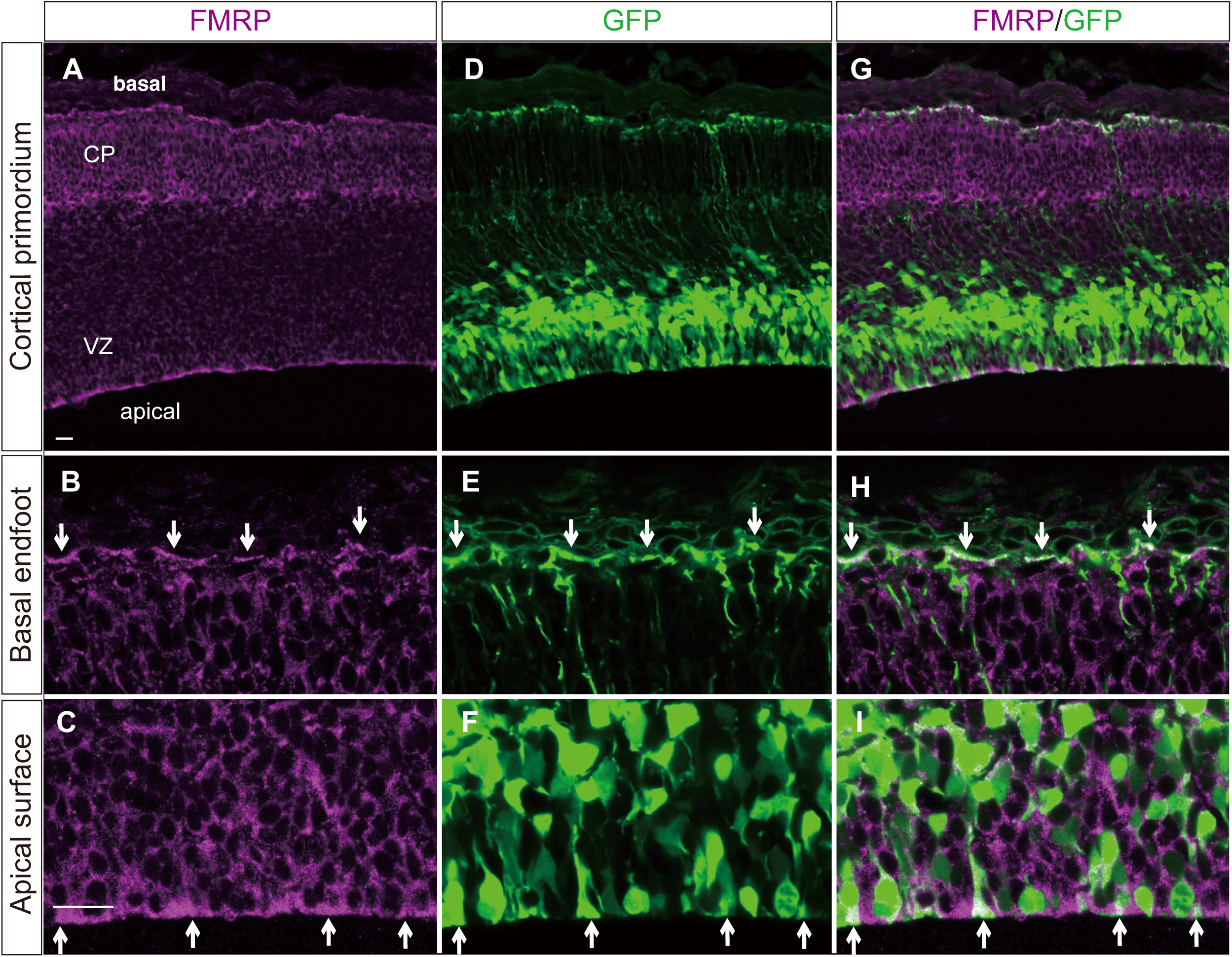
Expression patterns of FMRP in E14.5 WT brain. (A-C) Coronal sections of the neocortex showing the localization of FMRP at the basal endfeet and the apical surface of the RGCs (arrows). FMRP is also observed in the cortical plate (CP). (D-F) EGFP-labeled RGCs showing the basal and apical endfeet (arrows). (G-I) Merged images showing that FMRP is highly localized in the basal and apical endfeet of the RGCs. Scale bars represent 20 µm. VZ: ventricular zone

### Identification of FMRP target mRNAs in embryonic mouse cortex

To explore mRNA targets of FMRP during corticogensis, we performed RIP-sequencing (RIP-seq, Figure S1) using the cortical primodium samples isolated from E14.5 WT mice. In total, we found 3,954 transcripts that showed significant difference in expression (FDR<0.01), 2,288 of which were enriched in the FMRP-IP compared to the IgG-IP. We identified a stringent set of 947 transcripts from 865 candidate FMRP mRNA targets based on fold-change (log_2_FC>1) and FMRP-IP (FPKM>10) gene expression to prevent signal to noise issues from the sequencing (Figure 2A). This set of FMRP mRNA targets showed higher mRNA abundance compared to negative control and could thus be validated as candidate targets of FMRP in the developing neocortex.

**Figure 2.**
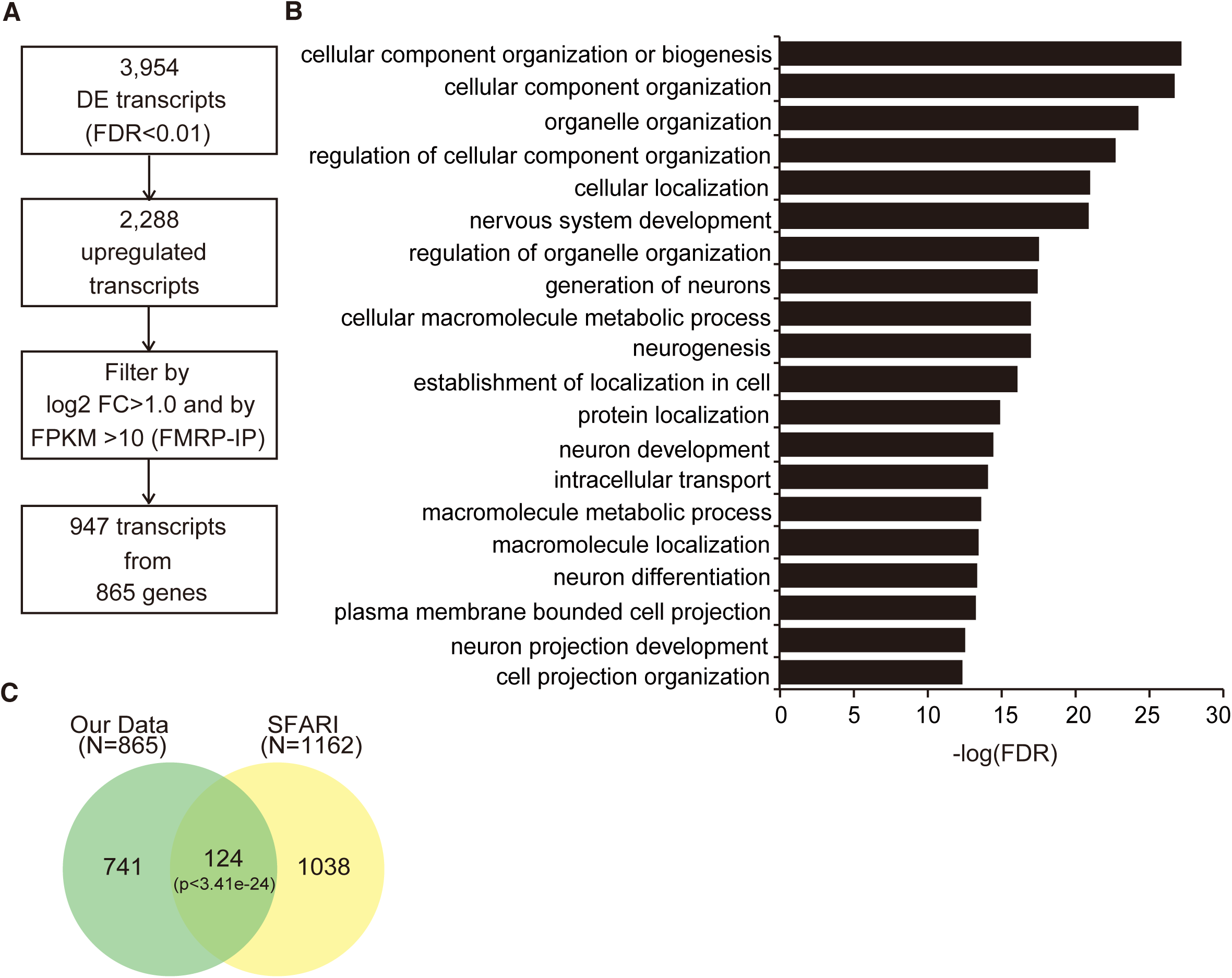
FMRP mRNA targets identified from the RIP-seq of WT E14.5 cortex. (A) Scheme used to identify FMRP mRNA targets from RIP-seq data. (B) The top 20 GO (biological process) terms enriched in FMRP targets genes by false discovery rate (FDR). (C) Venn diagram depicting the overlap of FMRP target genes identified in this study (green) and ASD-associated genes retrieved from the SFARI database (yellow). Statistically significant association was determined by hypergeometric distribution analysis.

To estimate functions of the FMRP target candidates, we used gene ontology (GO) term using the Visual Annotation Display (VLAD) – gene analysis and visualization analysis tool of the Mouse Genome Informatics (MGI) (Smith *et al.*, 2018). The top significant GO terms included biological processes involved in early brain development such as “nervous system development”, “generation of neurons”, “neurogenesis”, “neuron development”, “neuron differentiation”, and “neuron projection development” (Figure 2B). This is quite reasonable because FMRP has been shown to play an important role in maintaining the RGCs, i.e., the main source of neurons during brain development (Saffary and Xie, 2011; Götz and Barde, 2005). Thus, FMRP may participate in cortical neurogenesis via regulating various mRNA targets.

### Overlap the FMRP targets with ASD-associated genes

To search for a link between FXS to ASD, we compared our 865 FMRP target genes with 1162 ASD-associated genes from Simons Foundation Autism Research Initiative (SFARI, retrieved last March 11, 2019) (Abrahams *et al.*, 2013). There was a highly significant overlap of 124 candidate FMRP target genes and ASD-associated genes (p=3.41×10^−24^; Figure 2C, Supplementary Table 1). This overlap included several well-studied, syndromic ASD-associated genes like *HECT, UBA And WWE Domain Containing E3 Ubiquitin Protein Ligase 1* (*HUWE1*), *neurofibromatosis 1* (*NF1*), *Nipped-B homolog* (*NIPBL*), *nitric oxide synthase 1* (*NOS1*), *and paired box 6* (*PAX6*). The results obtained here may suggest a possible explanation for the common phenotypes of FXS and ASD during neural development.

### Protein-protein association network

To predict biological functions, pathways, and protein-protein interactions, 124 proteins encoded by both FMRP targets and ASD-associated genes were analyzed using the STRING Database (Szklarczyk *et al.*, 2017). Four statistically significant functional clusters were selected and highlighted (FDR<0.05; Figure 3). The cluster with the largest number of FMRP targets/ASD-associated genes was categorized as “transcription regulation” comprised of 28 genes, i.e., *Aff4, Arnt2, Atrx, Cic, Chd6, Chd8, Ctnnb1, Ep300, Kat6a, Kdm4b, Kdm6a, Kmt2c, Med12, Med13l, Myt1l, Nacc1, Ncor1, Nipbl, Nsd1, Pax6, Per1, Setd2, Setd5, Sin3a, Smarca4, Smarcc2, Taf1,* and *Tcf4* (see Supplementary Table 1 for full names of the genes). The second cluster was “regulation of actin cytoskeleton” comprised of 8 genes, i.e., *Apc, Cyfip1, Dock1, Fgfr2, Iqgap1, Mapk3, Nckap1,* and *Pik3r2*. The third cluster was “ubiquitin-mediated proteolysis” comprised of 7 genes, i.e., *Birc6, Cul7, Huwe1, Trip12, Ube2h, Ube3c* and *Ube3h.* The last cluster was “calcium signaling pathway” comprised of 5 genes, i.e., *Cacna1h, Gnas, Itpr1, Nos1* and *Plcb1.* Other proteins may also have the similar functions, although they did not appear in our current pathway analyses. Above evidence may imply multiple functions of FMRP through various target molecules.

**Figure 3.**
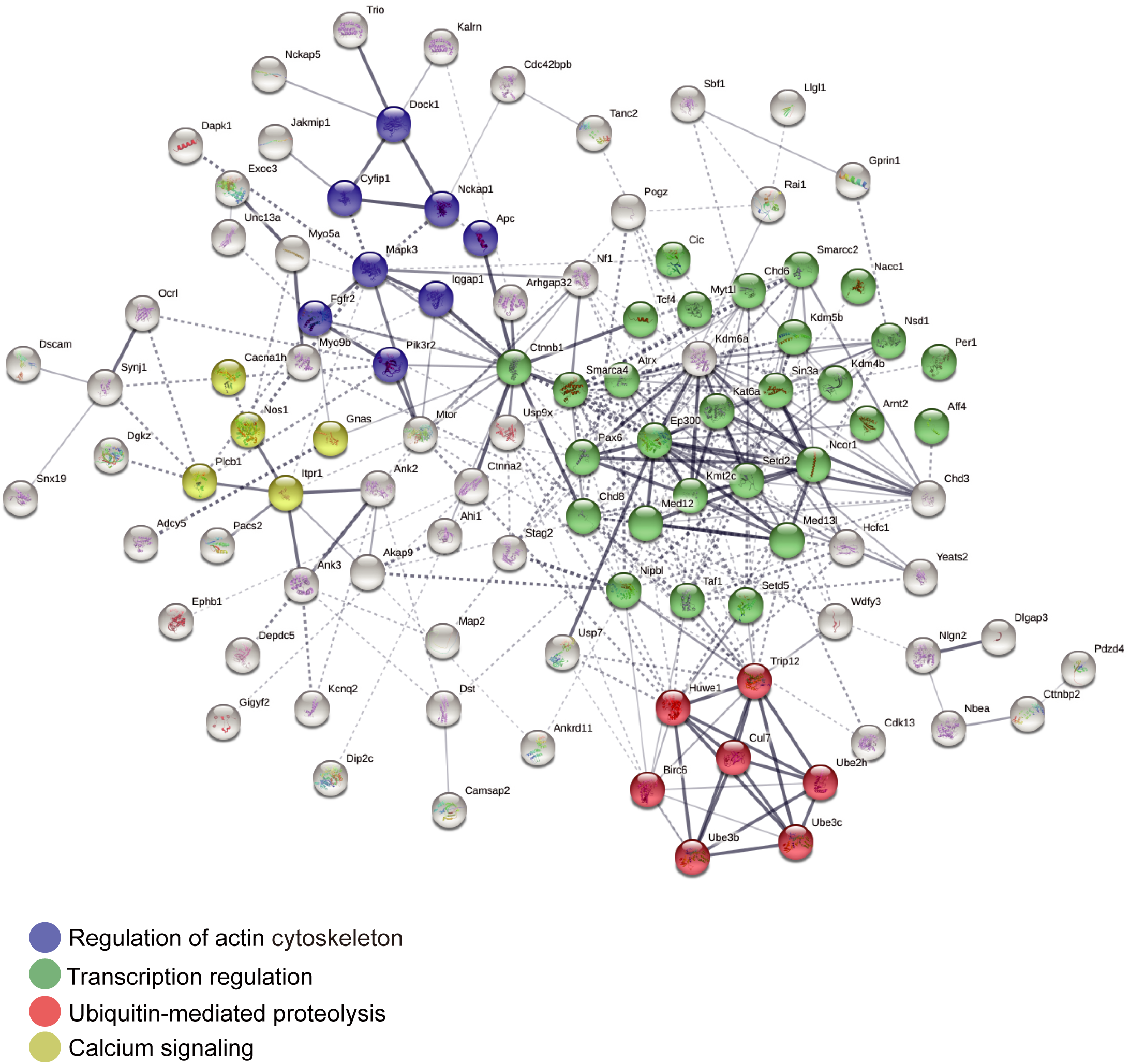
Protein-protein association network. Interaction of proteins encoded by the 124 overlapped genes analyzed by STRING database of predicted functional associations among genes/proteins. Stronger associations are represented by thicker lines. Four enriched statistically significant functions have been selected and highlighted: regulation of actin cytoskeleton (violet), transcription regulation (green), calcium signaling (yellow) and ubiquitin-mediated proteolysis (red).

### Confirmation of mRNA targets regulated by FMRP during corticogenesis

To validate the targets identified by RIP-seq, enrichment analysis of the mRNA targets in the four functional clusters was performed using RIP-qPCR of the embryonic dorsal telencephalon. 46 out of the 48 targets in the above mentioned four clusters showed significant enrichment in the FMRP-IP compared with that in the quality control (QC-input) (Figure 4). The top six most enriched targets with an expression fold-change greater than 20 (FC>20) were *Per1, Chd8, Cul7, Dock1, Pik3r2* and *Itpr1.* It is thus considered that the 46 targets are reliable as candidates for FMRP targets that may have impact on corticogenesis.

**Figure 4.**
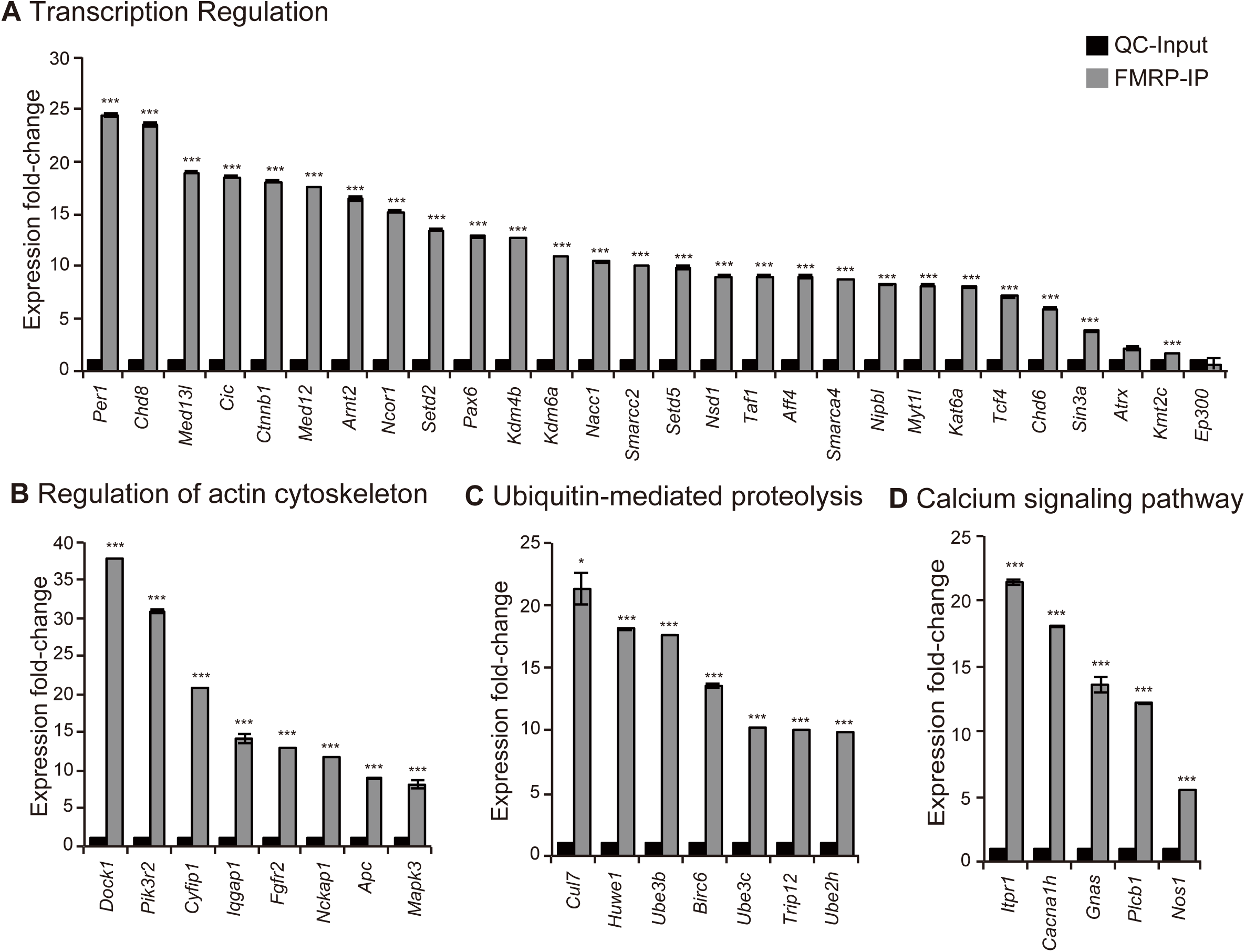
Validation of candidate FMRP targets by RIP-qPCR. The RNAs from WT (n=3) of the E15.5 mouse dorsal telencephalon were isolated and subjected to cDNA synthesis and qPCR. 46 out of 48 targets significantly enriched in the FMRP-IP compared with that of QC-input (n=3). Student’s *t-test* was used to test statistical significance (***p<0.0001).

The most known functions of FMRP are posttranscriptional regulation of its target mRNAs, e.g., transportation and translation within the cell (see review by Pilaz and Silver, 2015). However, we wondered if there is any possibility of regulation at the level of transcription. We thus performed RT-qPCR of FMRP target candidate genes using the embryonic dorsal telencephalon obtained from WT and *Fmr1-*KO mice at E15.5. Most of the 46 enriched FMRP targets did not show different expression levels between WT and *Fmr1-*KO mice, although there were four genes that showed difference in expression (Figure 5). *Huwe1* and *Kat6a* showed increased expression, while *Kmt2c* (also known as *Mll3*) and *Apc* showed decreased expression in *Fmr1-*KO mice. Therefore, it can be possible that these four mRNA targets are regulated at the level of transcription.

**Figure 5.**
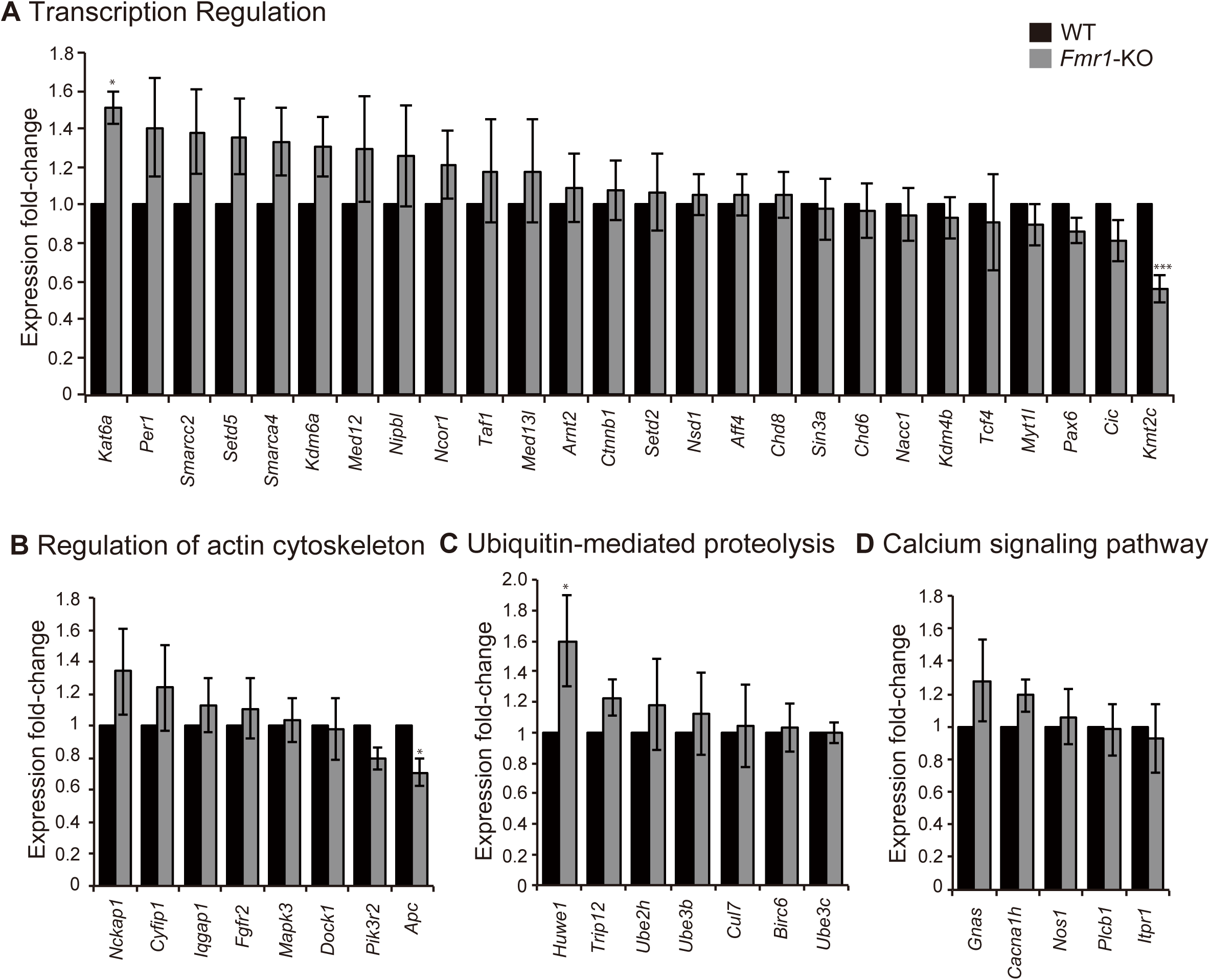
Expression of FMRP targets in the WT and *Fmr1*-KO mice. The RNAs from WT (n=7) and *Fmr1*-KO (n=7) of the E15.5 mouse dorsal telencephalon were isolated and subjected to cDNA synthesis and qPCR. Out of the 46 enriched mRNA targets, only 4 showed significant difference between the WT and *Fmr1-*KO. *Huwe1* and *Kat6a* showed increased expression while *Kmt2c* and *Apc* showed decreased expression in *Fmr1-*KO mice. Student’s *t-test* was used to test statistical significance (*p<0.01, ***p<0.0001).

### FMRP influences mRNA stability *Kmt2c* and *Apc*

To assess whether the decrease in mRNA amount of *Kmt2c* and *Apc in vivo* was resulted from reduced stability of mRNA or not, we performed mRNA stability assay using primary culture of cortical astrocytes and embryonic fibroblasts (MEFs) taken from *Fmr1*-KO and WT mice. We performed a time-course evaluation of mRNA amount of the genes in the cultured astrocytes and MEFs treated with actinomycin D, a transcription inhibitor (Bensaude, 2011). After transcriptional blockade, *Kmt2c* and *Apc* mRNA amount showed a similar level between WT and *Fmr1-*KO astrocytes (Figure 6A, B). In MEFs, mRNA amount of *Apc* was reduced by 13.5% in *Fmr1-*KO than that of WT at 4 hours after actinomycin D treatment, while *Kmt2c* mRNA showed reduction in mRNA amount in *Fmr1-*KO at 2-hr (26.3%) and 4 hours (40.9%) after the treatment (Figure 6C, D). These results suggest that FMRP could contribute to the mRNA stability of *Kmt2c*, at least, in cultured MEFs.

**Figure 6.**
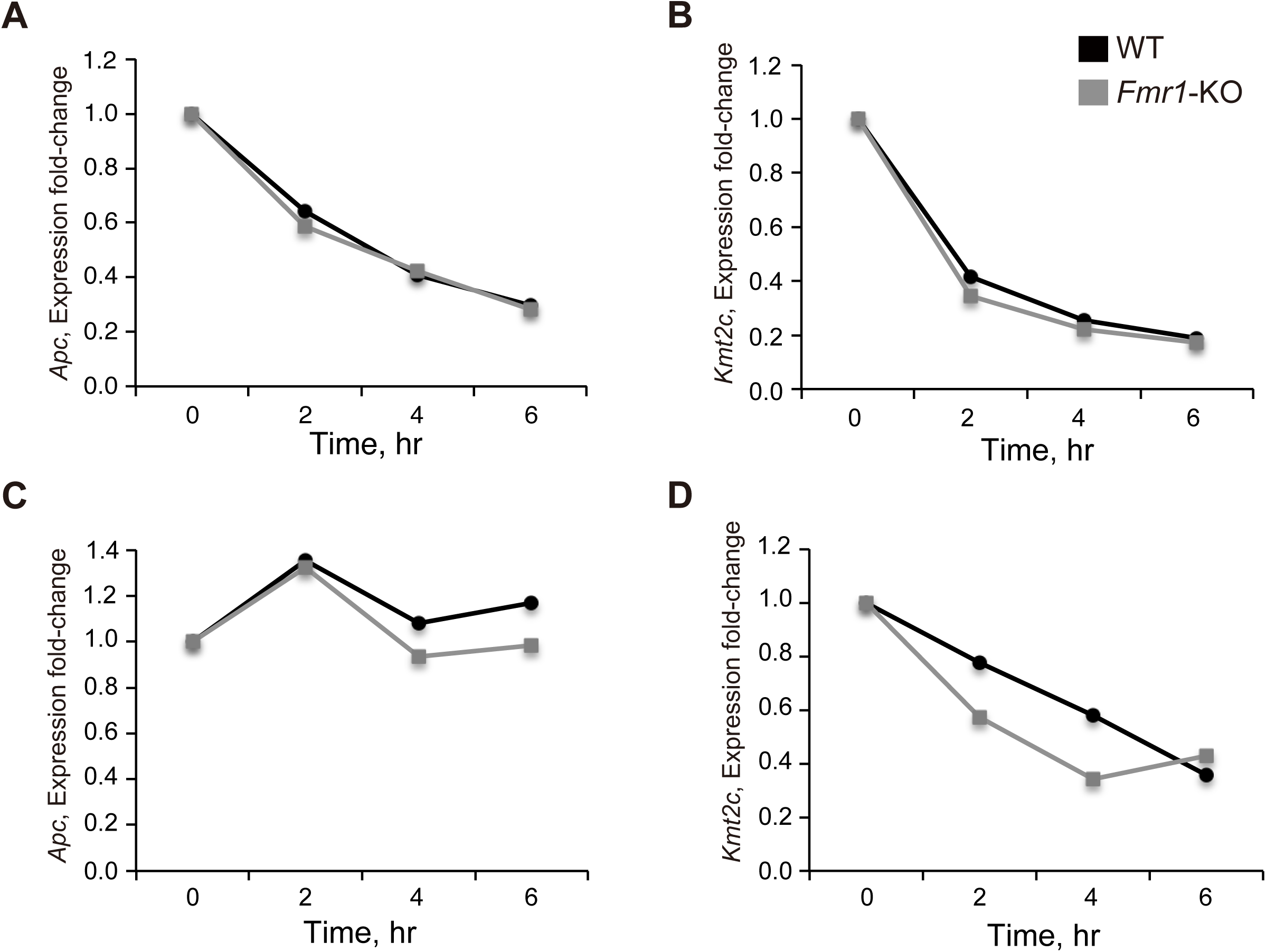
FMRP regulates the stability of *Kmt2c* mRNA in MEFs. Relative mRNA amount of the four genes in cultured cortical astrocytes (n=1) (A, B) and MEFs (n=1) (C, D) treated with actinomycin D at different time points. The stability of *Apc* mRNA was unaffected in both astrocytes and MEFs. *Kmt2c* mRNA was reduced in the MEFs from *Fmr1*-KO mice than that of WT, but unaffected in the astrocytes.

## DISCUSSION

Functions of FMRP have been reported to include regulation of translation, transport, and stability (De Rubeis and Bagni, 2010; Darnell *et al.*, 2011; Ascano *et al.*, 2012; La Fata *et al.*, 2014; Pilaz *et al.*, 2016; Liu *et al.*, 2018). Although mechanisms are not fully elucidated at molecular and cellular levels, several FMRP mRNA targets are linked to various neurobiological processes such as neuronal migration and circuitry formation (La Fata *et al.*, 2014), synaptogenesis (Darnell *et al.*, 2011), DNA damage response (Alpatov *et al.*, 2014), and epigenetic regulation (Korb *et al.*, 2017). Therefore, to identify mRNAs that are controlled by FMRP is essential for elucidation of its roles in brain development by controlling the target molecules and for determination of its contribution to pathogenesis of FXS and related neurodevelopmental disorders.

Here, we identified FMRP mRNA targets in murine corticogenesis. Among 865 genes obtained from RIP-seq, 124 are listed in SFARI database for autism-related genes, suggesting common roles of FMRP in FXS and ASD. In other words, various symptoms such as behavioral and cognitive deficits are commonly observed in both neurodevelopmental disorders possibly through such overlapped transcripts.

We categorized these common genes by protein network analyses into four clusters: transcription regulation, regulation of actin cytoskeleton, ubiquitin-mediated proteolysis, and calcium signaling pathway. Although previous studies on FMRP targets have already suggested a limited number of molecules related with above functional categories, our results clearly highlight importance of the whole molecules involved in these categories in corticogenesis.

The largest number of mRNA targets we found are genes whose products are classified into “transcription regulation” that controls the proliferation of neural stem cells and their transition to differentiating neural progenitors (Lee et al., 2014). One of the FMRP target in this category is *Pax6,* a key transcription factor in brain development by regulating hundreds of genes related to ASD (see review by Kikkawa et al., 2019). The mRNA and protein expression of *Pax6* in the *Fmr1-*KO mice showed a decreased tendency at below the significant level (Figure S2). The lack of the significance can be attributed to small sample size and/or weakness in the experimental design; or the regulation of Pax6 by FMRP may not be due to transcriptional regulation. A previous ChIP study has reported that *Fmr1* is one of the targets of Pax6 (Sansom *et al.*, 2009) and our preliminary work showed that FMRP expression was decreased in the *Pax6* deficient cortical primordium (data not shown). A positive feedback loop between these two pivotal molecules, i.e., FMRP and Pax6, could be important when we consider pathogenesis of neurodevelopmental disorders.

The second category we found is “regulation of actin cytoskeleton”. It is shown that FMRP regulates cell fate determination (Saffary and Xie, 2011), neuronal migration and cortical circuitry (La Fata *et al.*, 2014), and maintenance of the scaffold of RGCs (Yokota *et al.*, 2009) during brain development. Since these developmental processes are highly dependent on cytoskeletal organization, our mRNA targets of this category may play a role in these processes.

Another interesting group of genes that are considered to be ASD-related FMRP targets are related with “Ubiquitin-mediated proteolysis”. Ubiquitination is not only important in mature neurons (Hallengren *et al.*, 2013) but also in various neurodevelopmental processes such as synapse formation, neurogenesis, neurite enlargement, dendrite growth, axonal development, neural tube formation, and differentiation (Tuoc and Stoykova, 2010). Although ubiquitin-related genes, i.e., *Huwe1* and *Ube3b* are previously suggested as a target of FMRP in neurons (Darnell *et al.*, 2011), our data showing that FMRP can regulate 7 genes involved in the ubiquitin-mediated proteolysis may imply general importance of FMRP in ubiquitination.

Finally, we found ASD-related FMRP target genes categorized into “calcium signaling pathway”, although their number was only five. A previous literature has already mentioned molecules related with calcium signaling, a key process in neural functions (Davis and Broadie, 2018). This is quite expected because our original samples for RIP-seq were taken from the cortical primordium of E14.5, containing immature neurons where modulation of ion channels and regulation of excitability may occur. However, other possibility that calcium signaling is functioning within RGCs cannot be excluded.

In our study, we analyzed expression of ASD-related FMRP mRNA targets in the cortical primordium of *Fmr1-*KO by RT-qPCR. Only few studies have previously examined FMRP regulation on its targets at the mRNA level (Brown *et al.*, 2001; Korb *et al.*, 2017; Liu *et al.*, 2018). These reports do not directly demonstrate the mechanistic role of FMRP on the transcription of its target genes, but they provide clues that mRNA levels are altered if FMRP function is impaired.

Among the four ASD-related FMRP targets we found, *Kat6a* and *Kmt2c* are previously suggested to be FMRP mRNA targets from ribosomal profiling using adult neurons (Darnell *et al.*, 2011). Kat6a and Kmt2c are histone modification writers, which can epigenetically regulate transcription (Shen *et al.*, 2014; Tapias and Wang, 2017). Interestingly, expression of *Kat6a* and *Kmt2c* were conversely regulated by loss of FMRP functions. Though we did not demonstrate the interaction between *Kat6a* and *Kmt2c,* this result may suggest that a combinatorial or compensatory role of histone modifications may occur in the condition with FMRP dysfunction. Considering possible interaction of Kat6a and Kmt2c with chromatin, loss of FMRP could result in aberrant gene expression and widespread changes in chromatin regulation as mentioned by others (Alpatov *et al.*, 2014).

Another ASD-related FMRP target that showed decreased expression in *Fmr1-*KO mice, *Apc* has previously been described as a target of FMRP (Darnell *et al.*, 2011; Pilaz *et al.*, 2016), and loss of APC in the RGCs drastically disrupted RGC organization and inhibited appropriate proliferation of RGCs (Nakagawa *et al.*, 2017). In our data, *Apc* expression was decreased in *Fmr1-*KO cortical primordium, which may actually affect proliferation and organization of RGCs during brain development.

The last ASD-related FMRP target, Huwe1, showed increase in expression in loss of FMRP function. *Huwe1* is an a FMRP target in *Drosophila* oocytes, where decrease in *Huwe1* expression has been suggested due to indirectly enhanced translation by preventing protein degradation (Greenblatt and Spradling, 2018). The discrepancy can probably be accounted for by the function of FMRP in different tissues and that transcription of FMRP mRNA targets may also vary in different tissues (Zalfa *et al.*, 2007; Man *et al.*, 2017). Further studies on the regulation of FMRP on *Huwe1* during brain development might be relevant to understand its impact in the development of FXS.

The expression changes observed in *Kat6a, Kmt2c, Apc* and *Huwe1* were relatively small in the *Fmr1*-KO mice. Although, we do not know their functions during corticogenesis at this moment, subtle shifts in gene expression in some of the FMRP mRNA targets may collectively contribute directly or indirectly to the alteration of overall mRNA fates resulting in dysregulation of neurodevelopmental processes and subsequently in pathogenesis of FXS.

How does FMRP regulate transcription of its mRNA targets? In our study, *Kmt2c* showed reduced mRNA levels in *Fmr1-*KO embryonic brain, which might be due to loss of mRNA stability. Thus, we inferred that dysfunction of FMRP may reduce stabilization of *Kmt2c* mRNA, resulting in its decreased expression, which might contribute to impaired neural development in FXS.

Conversely, FMRP can also perturb gene expression like in cases of *Huwe1* and *Kat6a.* This can be direct, but also be indirect since it has been proven that FMRP interacts with multiple other RNA-binding proteins (Davis and Broadie, 2018). In case of a nuclear export factor NXF1, the absence of FMRP first inhibits NXF2 (also a nuclear export factor and close family of NXF1), which results in reduction of *Nxf1* expression (Zhang, et al,, 2007; Davis and Broadie, 2018). Changes of the gene expression levels of FMRP targets in the *Fmr1-*KO telencephalon could be regulated by FMRP directly or indirectly via other mechanisms i.e., epigenetic regulation and microRNAs.

In this study, we revealed that FMRP might interact with many mRNAs during mammalian corticogenesis. It is important to understand that not one single gene regulated by FMRP can explain a given phenotype of neurodevelopmental disorders. Instead, it may arise from the collective dysregulation of combinatory sets of FMRP targets. Though we do not know how FMRP mechanistically manages its mRNA targets, our findings suggest that dysfunctions of FMRP may influence changes the fate of its mRNA targets in various ways, multiple combination of which can reflect multifarious phenotypes of neurodevelopmental disorders.

## MATERIALS AND METHODS

### Animals

Animal experiments were carried out in accordance with the National Institutes of Health guidelines outlined in the Guide for the Care and Use Laboratory Animals. The Committee for Animal Experimentation of Tohoku University Graduate School of Medicine approved all the experimental procedures described herein (2017-MDA-189). Male WT (C57BL/6J) and *Fmr1*-KO mice (B6.129P2-*Fmr1*^*tm1Cgr*^/J, stock #003025) (Bakker *et al.*, 1994) were used in this study. Hemizygote (*Fmr1*^*-/y*^) male and heterozygote (*Fmr1*^*+/-*^) female mice were mated to obtain WT (*Fmr1*^*+/y*^) and *Fmr1-*KO (*Fmr1*^*-/y*^) male embryos.

### DNA extraction and genotyping

Deoxyribonucleic acid (DNA) was extracted from tail of E15.5 mouse embryos. A mixture of 10 µl of 5x Colorless GoTaq® Flexi Buffer (Promega). Standard polymerase chain reaction (PCR) was performed to determine gender and the WT and *Fmr1* KO alleles of the embryos using specific primers (Supplementary Table 2, gifted by Dr. Yukio Sasaki) and the GoTaq® Flexi DNA Polymerase (Promega). Standard PCR was performed using the Vapo Protect Thermal Cycler (Eppendorf), and the amplified PCR products were visualized by electrophoresis on 1% agarose gels using the Gel Doc™ EZ Imager (Bio-Rad).

### Immunohistochemistry

WT and *Fmr1*-KO embryos at E15.5 were perfused with 4% paraformaldehyde (PFA) in phosphate buffered saline (PBS), and the whole brain was collected. The embryonic brain was fixed in 4% PFA in PBS at 4°C for 2 hours, placed in 10% sucrose/PBS (weight/volume) overnight (8-12 hours), and in 20% sucrose/PBS for overnight or until the tissue sank. The embryonic brains were embedded in O.C.T. Compound (Sakura Finetek), frozen by dry ice, and stored at −80°C. Frozen coronal brain sections (14 µm) were prepared and were washed with Tris-buffered saline containing 0.1% Triton X-100 (TBST) for 5 minutes three times. The sections were placed into a moisture chamber (COSMO BIO), and incubated in 3% bovine serum albumin (BSA)/TBST for 1 hour at room temperature (RT). The sections were incubated with primary antibodies diluted with 3% BSA/TBST, including goat anti-FMRP (1:1000; LS-B3953; LifeSpan Biosciences Inc.) and rabbit anti-Pax6 (1:1000; Inoue et al. 2000) overnight at 4°C. Then sections were incubated with secondary antibodies diluted at 1:500 in 3%BSA/TBST for 1 hour at RT. The secondary antibodies used were Cy3-conjugated donkey anti-goat IgG (1:500, Life Technologies), Cy3-conjugated donkey anti-rabbit IgG (1:500, Life Technologies), and Alexa 488-conjugated donkey anti-mouse IgG (1:500). Cell nuclei were counterstained with 4’,6-Diamidino-2-phenylindole dihydrochloride (DAPI)/TBST (1:1000; Sigma). For observation, sections were mounted with VECTASHIELD® antifade mounting medium (Vector Laboratories Inc.), and sealed with cover slides by EUKIT (O.Kindler GmbH & CO). Images were visualized by a confocal laser microscope Zeiss LSM780 (Carl Zeiss). The signals were quantified using ImageJ 1.48v software (NIH). Corrected total cell fluorescence was computed by measuring the fluorescence intensity of target in 40 cells and intensity of the background (40 areas).

### *In utero electroporation* into mouse embryonic brain

*In utero* electroporation was performed as described previously with minor modification (Saito and Nakatsuji, 2001; Sato *et al.*, 2013). Pregnant WT mice at E13.5 were anesthetized with by isoflurane for laboratory animals (MSD Animal Health) using the Univentor 400 anaesthesia unit and vaporizer (Univentor). For surgery, the uterus of the pregnant mouse was exposed after making approximately 3 cm of incision in the middle abdominal region. The expression vectors pCAG-EGFP plasmid (kindly gifted from Prof. Tetsuichiro Saito, Chiba University, Japan) and 1% Fast green in PBS was injected into the lateral ventricle of embryos at E13.5 with a mouth-controlled glass capillary pipette. Immediately, square pulses (40 V, 50 ms, five times at 1 second intervals) were delivered into embryos with an electroporator (CUY-21, BEX) and a forceps-type electrode (LF650P5, BEX). After the electroporation, the uterus was returned back to inside of the abdomen and sutured with Suture with Needle kit (8 mm of diameter curved needle, 45 cm of thread, BEAR Medic Corporation). Finally, the most out layer of skin was clipped by 9 mm stainless steel AUTOCLIP Applier (Becton Dickinson and company). After surgery, mice were placed on a 37°C heating pad for recovery. Embryos were collected at E14.5 for further analysis.

### Preparation of RNA libraries and sequencing

Following the manufacturer’s protocol, RIP assay was performed to extract FMRP-bound mRNAs from the dorsal telencephalon of WT mice at E14.5 by using RiboCluster Profiler™ RIP-Assay Kit, anti-FMRP Human polyclonal antibody, #RN016P, and Dynabeads™ Protein beads G/A (Invitrogen™). The quality and quantity of the total RNA was evaluated using the Agilent 2100 Bioanalyzer with RNA 6000 Pico Kit (Agilent). Total RNA concentration greater than 50 ng with an RNA Integrity Number (RIN) value greater than or equal to 7.9 (≥7.9) were submitted to the Graduate School of Frontier Sciences, The University of Tokyo for next generation sequencing. A total of three cortices immunoprecipitated with anti-FMRP and two cortices immunoprecipitated with anti-IgG were sequenced.

### Sequence alignment and estimation of gene expression levels

To trim the raw reads containing the Illumina adaptors and remove low quality sequences, the FASTX-Toolkit (http://hannonlab.cshl.edu/fastx_toolkit/) was used. Quality reads below 20 and sequence length below 70 were discarded. Only good quality and paired reads were analyzed in the next step.

TopHat-Cuffdiff pipeline was employed to analyze the RNA sequences (http://tophat.cbcb.umd.edu/, http://cufflinks.cbcb.umd.edu/). Using TopHat, sequence reads were aligned to the *Mus musculus* genome (mm10) with default parameter values, except for distance between mature pairs (*r*=200), or the gap between paired reads. The final efficiency of alignment of all the sequences was between 72 to 87% (Supplementary Table 3). Gene expression levels were calculated as fragment per kilobase of transcript per RNA-seq read mapped (FPKM) using Cuffdiff. We defined the enriched RNAs after FMRP pulldown as FMRP-IP and the negative control as IgG-IP. The fold change (FC) on a logarithmic scale with base 2 (log_2_FC) and *p-*value for each gene are calculated to statistically test differential expression between the two conditions (FMRP-IP versus IgG-IP). We only used genes showing nominally significant difference at Q<0.01, fold-change equal to or greater than 1, and FPKM value greater than 10 in the FMRP-IP samples in the subsequent analyses of differentially expressed genes (DEGs) (Figure 2B).

### Gene Ontology and Protein Association Network

Functional annotation of the DEGs was performed using the VLAD tool (v1.6.0) of the MGI. GO was determined *via* an enrichment analysis (biological process) and the false discovery rate (FDR) less than 0.05 was considered as significantly enriched GO annotation (Supplementary Table 4). Protein network analysis was performed using the STRING database of known and predicted protein-protein interactions (Szklarczyk *et al.*, 2017).

### RNA extraction, quantitative PCR and RIP-qPCR

Total RNA was isolated with the RNeasy Mini Kit (Qiagen) according to the manufacturer’s protocol. RIP-qPCR was performed to validate and quantify expression of the FMRP targets using the FMRP-IP and QC-input dorsal telencephalon samples (pooled from WT, n=3), and dorsal telencephalon samples from WT (n=7) and *Fmr1-*KO (n=7) mice, respectively, for RT-qPCR. The complimentary DNA (cDNA) was synthesized using SuperScipt™III first-Strand Synthesis System for RT-PCR (Invitrogen). qPCR was performed Mastercycler® ep Gradient Realplex 2 (Eppendorf) with 2x SsoAdvanced Universal SYBR®Green Supermix (Roche) according to the manufacturer’s protocol. The threshold cycle (C_t_) values were obtained and fold change of expression (ΔΔCt) was calculated with *Rplp0* as normalizer. PCR sequences for qPCR (Supplementary Table 5) were designed by PrimerBank (https://pga.mgh.harvard.edu/primerbank/) and from previous reports (Duong *et al.*, 2011; Bonnans *et al.*, 2012; Armoskus *et al.*, 2014; Korneev *et al.*, 2015; Durak *et al.*, 2016; Yasuma *et al.*, 2016; Baell *et al.*, 2018; Cheng *et al.*, 2019).

### Western blot

Protein samples were collected from the pooled (n=3) dorsal telencephalon of WT and *Fmr1-*KO embryos at E15.5. The samples were lysed with cell lysis buffer (1 M HEPES pH 7.5, 80% glycerol, 5 M NaCl, 1 M MgCl_2_, 1 M DTT, 0.1 M PMSF, 10% NP40, X100 protease and phosphatase inhibitor, 0.5 M EDTA) on ice, and homogenized with a homogenizer (Nippi, Incorporated). Lysates were sonicated for 4 cycles of 30 seconds, and separated by centrifugation at 4°C and 15,000 rpm for 10 minutes. Protein concentration was measured by NanoDrop One/One Spectrophotometer (Thermo Scientific). The protein samples were heated in sodium dodecyl sulfate (SDS) gel loading buffer (1 M Tris pH 6.8, 1 M DTT, SDS, 10% BPB, glycerol) for 10 minutes before loading onto 6% SDS-polyacrylamide electrophoresis (SDS-PAGE gel) at 50 µg per lane. After running the gel, proteins were transferred electrophoretically onto polyvinylidene difluoride (PVDF) membranes (Millipore) with 40 V at 4°C for 720 minutes. The membranes were then blocked in 10% blocking buffer (Licor) in PBS for 1 hour, and incubated with a primary antibody, either goat anti-FMRP (1:1000; BD LSB3952) or rabbit anti-Pax6 (1:1000; Inoue et al. 2000), diluted in 10% blocking buffer with PBS at 4°C overnight. Anti-goat GAPDH (1:1000; Abcam) was used as loading control. The membrane was washed with TBST (containing 0.1% Tween 20) for 1 minute with three repeats and for 5 minutes with three repeats, and incubated with a secondary antibody, either of donkey anti-goat 800 (1:10000; Licor), goat anti-rabbit 680 (1:10000; Licor), or donkey anti-mouse 680 (1:10000; Licor), diluted in 10% blocking buffer with PBS for 1 hour at RT under a shaded condition. The fluorescence was detected using ODYSSEY infrared imaging system (Licor). The signals were quantified using ImageJ 1.48v software (NIH).

### *In situ* hybridization

*In situ* hybridization was performed as previously described (Osumi *et al.*, 1997; Kikkawa *et al.*, 2013). Frozen sections were incubated in Proteinase K (0.3 µg/ml) in PBS containing 0.1% Tween 20 (PBST) in 37°C water bath for 8 minutes. The sections were washed twice with PBST for 1 minute, and at RT for 2 minutes. The sections were then incubated in 4% PFA for 20 minutes. Ten-µl of RNA probe (50 ng/ml) was mixed with 150 µl of hybridization buffer (HB:100% Formamid, 20x SSC, 10% SDS, 50 µg/ml Heparin, 50 µg/ml tRNA from *Escherichia coli* (*E. coli*), diluted in RNase-free water). The *Pax6* RNA probe was produced by cloning the WT cDNA fragment amplified with the following primers: 5’-TCAGCTTGGTGGTGTCTTTG-3’ (sense) and 5’-GGTTGCATAGGCAGGTTGTT-3’ (antisense). The amplified sequences (inserts) were ligated to Bluescript vector (Promega) and transformed in competent cells (*E. coli* DH5α). RNA probes were then synthesized using polymerases T3/T7 (Promega). The probes in HB were applied on the sections, covered with coverslip (GRACE Biolab), placed into a moisture chamber (COSMO BIO), and then incubated at 70°C for 14-16 hours.

The sections were washed in Solution I (100% Formamid, 20x SSC, 10% SDS, milliQ water) at 67.5°C for 30 minutes, twice with Solution III (100% Formamid, 20x SSC, milliQ water) at 67.5°C for 45 minutes, thrice with Tris-buffered saline containing 0.1% Tween-20 (TBST) at RT for 5 minutes, and followed by blocking with 0.5% Blocking Reagent in PBST at RT for 1 hour. The sections were incubated with anti-Digoxigenin-AP Fab fragments (α-DIG, 1:4000, Roche), placed in a moisture chamber, and incubated overnight at 4°C. For color reaction, the section were washed thrice with TBST for 20 min at RT, followed by NTMT solution (5M NaCl, 1M MgCl_2_, 2M Tris-HCL, 10% Tween 20, milliQ water) at RT for 5 minutes and then incubated in the color reaction reagent (NTMT solution, 50 mg/ml NBT, 50 mg/ml BCIP) until the signal started to show up (approximately 8-24 hours). The sections were mounted with VECTASHIELD mounting medium, and sealed with cover slides by EUKIT. Images were visualized by a bright field microscopy (Keyence BZ-X710 All-in-One Fluorescence Microscope).

### Astrocyte culture

Astrocytes were cultured following essentially our study (Sakurai and Osumi, 2008). WT and *Fmr1-*KO pups at postnatal day 2 (P2) were anesthesized deeply and sterilized by dipping in 70% ethanol. Lateral cortices were dissected from hemispheres in 2 ml of ice-cold Tyrode’s solution and dispersed mechanically in 1 ml of Dulbecco’s modified eagle medium (DMEM) containing 10% fetal bovine serum (Sigma), 1% GlutaMAX (Gibco), and 1% antibiotic-antimycotic (Gibco). The dispersed cells were collected by centrifugation at 1000 rpm for 5 minutes, resuspended in 1 ml of medium, transferred into T75 flask with 10 ml of media, and cultured in the 37°C incubator (5% CO_2_) for 9 days. Media was exchanged every 3 days. At day 9, the cultured astrocytes were rinsed three times with 5 ml of PBS and once with 5 ml of media. Ten-ml of fresh media was added into T75 flask and the flasks were incubated for over 2 hours to stabilize pH of media. The flasks were then shaken vigorously at 250-300 rpm in the 37°C incubator for 18-24 hours to remove contaminated neurons, oligodendrocytes, and microglia. After shaking, cells were rinsed cells with 5 ml of PBS three times and incubated with 2 ml of trypsin-EDTA (Gibco) for 5 minutes. The enzymatic reaction was stopped by adding 8 ml of media, cells were collected by centrifugation at 1000 rpm for 5 minutes. Cell pellets were resuspended in the media and resuspended cells were seeded at 10^6^ cells/dish in 3.5-cm dishes. The primary astrocytes were cultured for 3 days and used for mRNA stability assay.

### Mouse embryonic fibroblast culture

The protocol was according to that used in a previous study with minor modification (Durkin *et al.*, 2016). MEFs were derived from WT and *Fmr1-*KO E14.5 embryos. The head, internal organs and limbs were removed from the embryos, and the trunk was mechanically minced and transferred in a 50 ml tube with 10 ml in Hank’s calcium magnesium free (HCMF) solution. HCMF solution was removed and replaced with 9 ml 0.25% trypsin-EDTA (Gibco), and samples were incubated at 37°C for 15 minutes. To stop enzymatic reaction, 10 ml of DMEM was added and then the tubes centrifuged at 1000 rpm for 5 minutes. The sticky material and the supernatant were removed from the samples using a pipette. The remaining cells were re-suspended in in 10-mm plate containing 10 ml DMEM, containing 10% fetal bovine serum (Sigma), 1% GlutaMAX (Gibco), and 1% antibiotic-antimycotic (Gibco) and incubate in the 37°C incubator (5% CO_2_). Cells were cultured for 9 days and used for mRNA stability assay.

### mRNA stability assay

For blocking transcription, both cultured MEFs and astrocytes were incubated with 1 ml of media containing 10 µg/ml of actinomycin D (dissolve in dimethyl sulfoxide, DMSO) (Wako) following a previous protocol (Zalfa *et al.*, 2007) After treatment for 2, 4 and 6 hours, the treated and untreated (DMEM and with 0.001% DMSO) cells were rinsed once with 1 ml of PBS, and collected with 1 ml of TRIzol (Invitrogen). RNA was extracted as following the instruction manual of TRIzol and further purified with RNeasy Mini Kit (Qiagen). Purified RNA was used for RT-qPCR.

### Statistical analysis

Data were compiled using Microsoft Excel 2011 and Student’s *t*-test was used to calculate statistical significance. Hypergeometric distribution was calculated using the webtool Hypergeometric Distribution Calculator (https://keisan.casio.com/exec/system/1180573201). Values of p<0.05 were considered significant.

## ACKNOWLEDGEMENT

We thank Dr. Yutaka Suzuki, Dr. Yuta Kuze, Ms. Kiyomi Imamura (Laboratory of Systems Genomics, Department of Computational Biology and Medical Sciences, Graduate School of Frontier Sciences, The University of Tokyo) and Dr. Sumio Sugano (Department of Medical Genome Sciences, Graduate School of Frontier Sciences, The University of Tokyo) for sequencing using the NGS. We are grateful to Dr. Yoshio Wakamatsu (Department of Developmental Neuroscience, Tohoku University Graduate School of Medicine) for sharing his cloning techniques and fruitful discussion, to Dr. Tatsuya Sato (Frontier Research Institute for Interdisciplinary Sciences, Tohoku University) for the electroporation technique, and to Dr. Yukio Sasaki (Functional Structure Biology Laboratory, Division of Functional Structure, Yokohama City University) for initially providing *Fmr1-*KO mice and genotyping primers.

## AUTHOR CONTRIBUTIONS

Conceptualization: C.C., T.K., H.I., N.O.; Methodology: C.C., H.I.; Analysis: C.C; Resources: C.C., T.K., H.I., N.O.; Data curation: C.C., H.I.; Data analysis: C.C.; Writing-draft: C.C.; Writing-review and editing: C.C., T.K., H.I., N.O.; Project administration: T.K.; Funding: T.K., N.O.

## FIGURE LEGENDS

**Figure S1.**
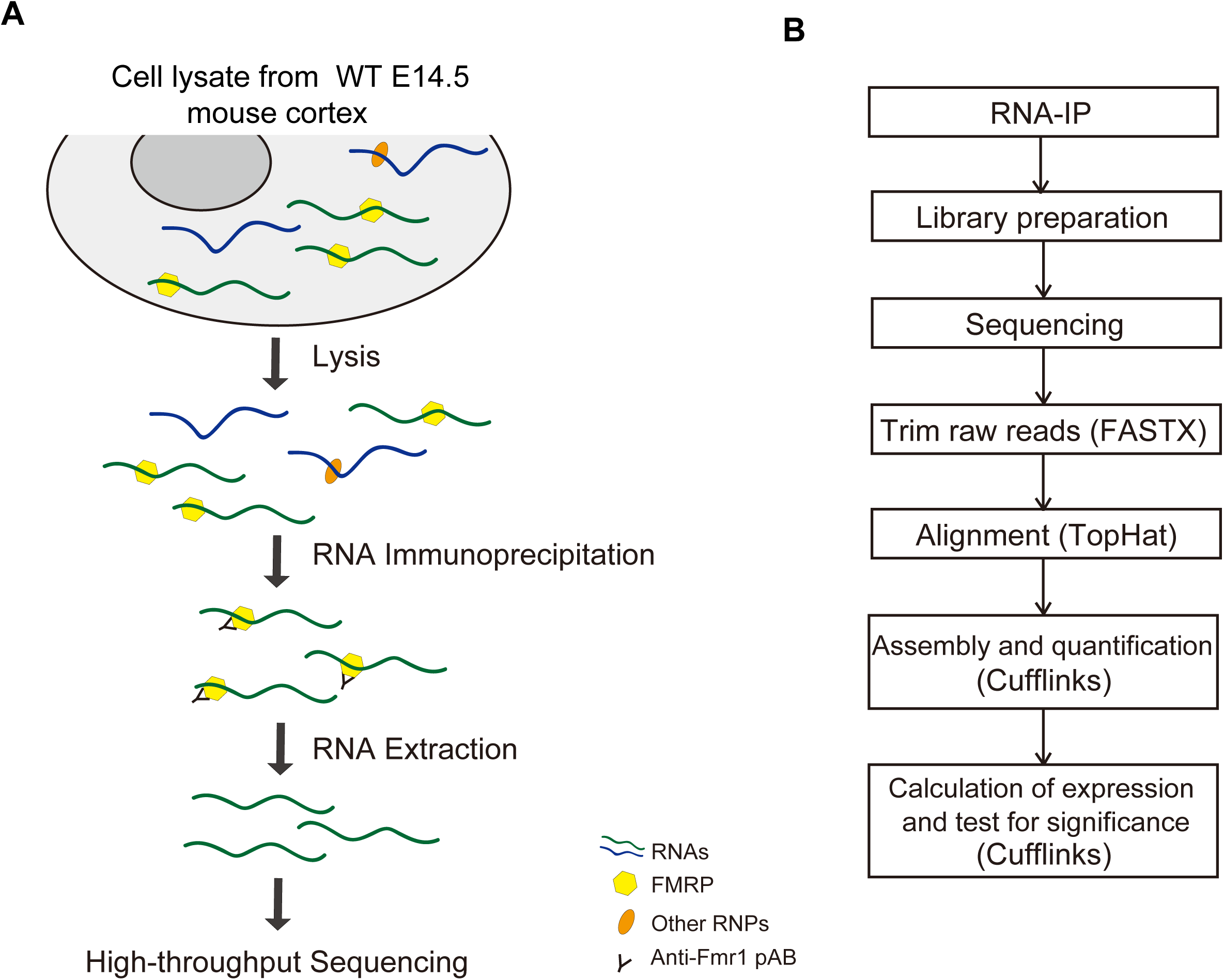
RNA immunoprecipitation sequencing (RIP-seq) workflow. (A) Summary of the RIP assay. (B) Outline of the RIP-seq data analysis.

**Figure S2.**
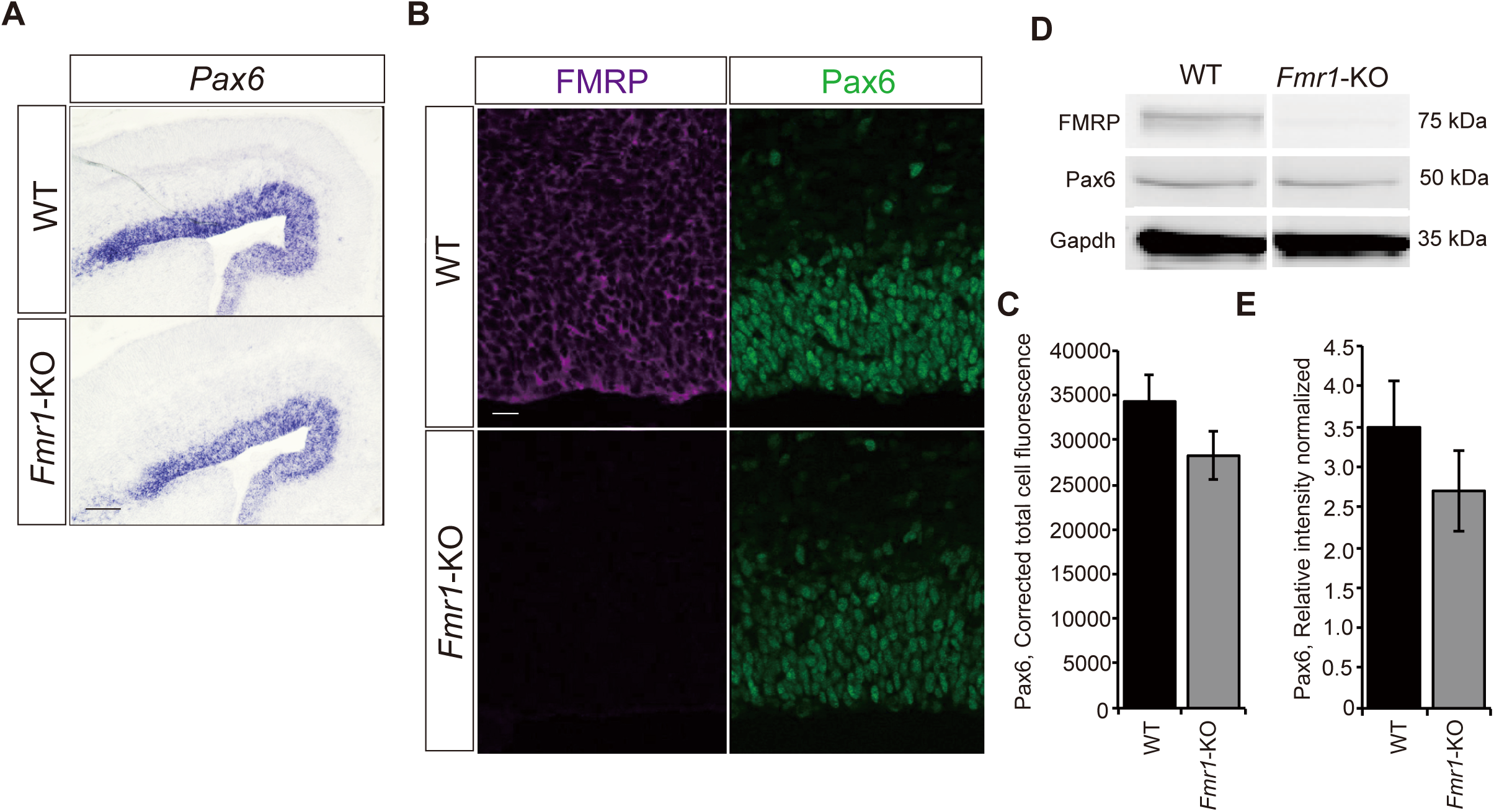
*Pax6* is one of the target genes identified in this study. (A) *In situ* hybridization (ISH) (n=3) showing a slight decrease in *Pax6* expression in the *Fmr1-*KO mice. (B-E) From the immunostaining (n=5) and western blot (n=6) results, though not statistically significant, there seemed to be a decrease in Pax6 expression in *Fmr1-*KO. Samples analyzed were all male, E15.5 dorsal telencephalon of the WT and *Fmr1-*KO embryos. The error bars reflect the standard error of the mean (SEM). Scale bars represent 100 µm.

## LITERATURE CITED

Abrahams, B. S. et al. (2013) ‘SFARI Gene 2.0: a community-driven knowledgebase for the autism spectrum disorders (ASDs)’, Molecular Autism, (4), p. 36.

Abrahams, B. S. and Geschwind, D. H. (2008) ‘Advances in autism genetics: On the threshold of a new neurobiology’, Nature Reviews Genetics, 9(5), pp. 341–355.

Alpatov, R. et al. (2014) ‘A chromatin-dependent role of the fragile X mental retardation protein FMRP in the DNA damage response’, Cell, 157(4), pp. 869–881.

Armoskus, C. et al. (2014) ‘Identification of sexually dimorphic genes in the neonatal mouse cortex and hippocampus’, Brain Research, 1562, pp. 23–38.

Ascano, M. et al. (2012) ‘FMRP targets distinct mRNA sequence elements to regulate protein expression’, Nature, 492(7429), pp. 382–386.

Ashley Jr., C. T. et al. (1993) ‘FMR1 protein?: Conserved RNP family domains and selective RNA binding’, Science, 262(October), pp. 563–567.

Baell, J. B. et al. (2018) ‘Inhibitors of histone acetyltransferases KAT6A/B induce senescence and arrest tumour growth’, Nature, 560(7717), pp. 253–257.

Bakker, C. E. et al. (1994) ‘Fmrl Knockout M ice? A Model to Study Fragile X Mental Retardation’, Cell, 78, pp. 23–33.

Bassell, G. J. and Warren, S. T. (2008) ‘Fragile X syndrome: Loss of local mRNA regulation alters synaptic development and function’, Neuron, 60(2), pp. 201–214.

Bensaude, O. (2011) ‘Inhibiting eukaryotic transcription: Which compound to choose? How to evaluate its activity?’, Transcription, 2(3), pp. 103–108.

Bonnans, C. et al. (2012) ‘Essential requirement for β-arrestin2 in mouse intestinal tumors with elevated Wnt signaling’, Proceedings of the National Academy of Sciences, 109(8), pp. 3047–3052.

Brown, V. et al. (2001) ‘Microarray Identification of FMRP-Associated Brain mRNAs and Altered mRNA Translational Profiles in Fragile X Syndrome’, Cell, 107, p. 27710.

Castrén, M. L. (2016) ‘Cortical neurogenesis in fragile X syndrome.’, Frontiers in Bioscience (Scholar edition), 8, pp. 160–168.

Cheng, Y. et al. (2019) ‘KDM4B protects against obesity and metabolic dysfunction’, Proceedings of the National Academy of Sciences, 116(6), pp. 2384–2385.

Crawford, D. C. et al. (2001) ‘FMR1 and the fragile X syndrome: Human genome epidemiology review’, Genetics in Medicine, 3(5), pp. 359–371.

Dahlhaus, R. (2018) ‘Of men and mice: Modeling the fragile X syndrome’, Frontiers in Molecular Neuroscience, 11(March), pp. 1–38.

Darnell, J. C. et al. (2011) ‘FMRP stalls ribosomal translocation on mRNAs linked to synaptic function and autism’, Cell, 146(2), pp. 247–261.

Davis, J. K. and Broadie, K. (2018) ‘Multifarious functions of the Fragile X mental petardation protein’, Trends in Genetics, 33(10), pp. 703–714.

Duong, H. A. et al. (2011) ‘A molecular mechanism for circadian clock negative feedback’, Science, 332(6036), pp. 1436–1439.

Durak, O. et al. (2016) ‘Chd8 mediates cortical neurogenesis via transcriptional regulation of cell cycle and Wnt signaling’, Nature Neuroscience, 19, p. 1477.

Durkin, M. E. et al. (2016) ‘Isolation of Mouse Embryo Fibroblasts Materials and Reagents’, 3(18).

La Fata, G. et al. (2014) ‘FMRP regulates multipolar to bipolar transition affecting neuronal migration and cortical circuitry’, Nature Neuroscience, 17(12), pp. 1693–1700.

Garber, K., Visootsaka, J. and Warren, S. (2008) ‘Fragile X syndrome’, European Journal of Human Genetics, 16, pp. 666–672.

Götz, M. and Barde, Y. A. (2005) ‘Radial glial cells: Defined and major intermediates between embryonicstem cells and CNS neurons’, Neuron, 46(3), pp. 369–372.

Greenblatt, E. J. and Spradling, A. C. (2018) ‘Fragile X mental retardation 1 gene enhances the translation of large autism-related proteins’, Science, 361(August), pp. 709–712.

Hallengren, J., Chen, P.-C. and Wilson, S. M. (2013) ‘Neuronal ubiquitin homeostasis’, Cell Biochemistry and Biophysics, 67(1), pp. 67–73.

Kikkawa, T. et al. (2013) ‘Dmrta1 regulates proneural gene expression downstream of Pax6 in the mammalian telencephalon’, Genes to Cells, 18(8), pp. 636–649.

Kikkawa, T. et al. (2019) ‘The role of Pax6 in brain development and its impact on pathogenesis of autism spectrum disorder’, Brain Research, 1705, pp. 95–103.

Korb, E. et al. (2017) ‘Excess translation of epigenetic regulators contributes to fragile X syndrome and Is alleviated by Brd4 inhibition’, Cell, 170(6), pp. 1209-1223.e20.

Korneev, S. A. et al. (2015) ‘A novel long non-coding natural antisense RNA is a negative regulator of Nos1 gene expression’, Scientific Reports. 5, p. 11815.

Lee, H.-K., Lee, H.-S. and Moody, S. A. (2014) ‘Neural Transcription Factors: from Embryos to Neural Stem Cells’, Molecules and Cells, 37(10), pp. 705–712.

Liu, B. et al. (2018) ‘Regulatory discrimination of mRNAs by FMRP controls mouse adult neural stem cell differentiation’, Proceedings of the National Academy of Sciences, 115(48), pp. E11397–E11405.

Man, L. et al. (2017) ‘Fragile X-associated diminished ovarian reserve and primary ovarian insufficiency from molecular mechanisms to clinical manifestations’, Frontiers in Molecular Neuroscience, 10(September), pp. 1–17.

Maurin, T. et al. (2018) ‘HITS-CLIP in various brain areas reveals new targets and new modalities of RNA binding by fragile X mental’, Nuclear Acids Research, 46(12), pp. 6344–6355.

Nakagawa, N. et al. (2017) ‘APC sets the Wnt tone necessary for cerebral cortical progenitor development’, Genes and Development, 31, pp. 1679–1692.

Osumi, N. et al. (1997) ‘Pax-6 is involved in the specification of hindbrain motor neuron subtype’, Development, 124(15), pp. 2961–2972.

Pilaz, L. J. et al. (2016) ‘Dynamic mRNA transport and local translation in radial glial progenitors of the developing brain’, Current Biology, 26(24), pp. 3383–3392.

Pilaz, L. and Silver, D. L. (2015) ‘Post-transcriptional regulation in corticogenesis?: how RNA-binding proteins help build the brain’, Wiley Interdisciplinary Reviews: RNA, 6(October), pp. 501–515.

De Rubeis, S. and Bagni, C. (2010) ‘Fragile X mental retardation protein control of neuronal mRNA metabolism: Insights into mRNA stability’, Molecular and Cellular Neuroscience, 43(1), pp. 43–50.

Saffary, R. and Xie, Z. (2011) ‘FMRP regulates the transition from radial glial cells to intermediate progenitor cells during neocortical development’, Journal of Neuroscience, 31(4), pp. 1427–1439.

Saito, T. and Nakatsuji, N. (2001) ‘Efficient Gene Transfer into the Embryonic Mouse Brain Using in Vivo Electroporation’, Developmental Biology, 246, pp. 237–246.

Sakurai, K. and Osumi, N. (2008) ‘The neurogenesis-controlling factor, Pax6, inhibits proliferation and promotes maturation in murine astrocytes’, Journal of Neuroscience, 28(18), pp. 4604–4612.

Sansom, S. N. et al. (2009) ‘The level of the transcription factor Pax6 is essential for controlling the balance between neural stem cell self-renewal and neurogenesis’, PLoS Genetics, 5(6), pp. 20–23.

Sato, T., Muroyama, Y. and Saito, T. (2013) ‘Inducible gene expression in postmitotic neurons by an in vivo electroporation-based tetracycline system’, Journal of Neuroscience Methods, 214(2), pp. 170–176.

Shen, E. et al. (2014) ‘Regulation of histone H3K4 methylation in brain development and disease’, Philosophical Transactions of the Royal Society B: Biological Sciences, 369(1652).

Smith, C. L. et al. (2018) ‘Mouse Genome Database (MGD)-2018: Knowledgebase for the laboratory mouse’, Nucleic Acids Research, 46(D1), pp. D836–D842.

Szklarczyk, D. et al. (2017) ‘The STRING database in 2017: Quality-controlled protein-protein association networks, made broadly accessible’, Nucleic Acids Research, 45(D1), pp. D362–D368.

Tapias, A. and Wang, Z. Q. (2017) ‘Lysine Acetylation and Deacetylation in Brain Development and Neuropathies’, Genomics, Proteomics and Bioinformatics, 15(1), pp. 19–36.

Tuoc, T. C. and Stoykova, A. (2010) ‘Roles of the ubiquitin-proteosome system in neurogenesis’, Cell Cycle, 9(16), pp. 3174–3180.

Verkerk, A. J. M. H. et al. (1991) ‘Identification of a gene (FMR-1) containing a CGG repeat coincident with a breakpoint cluster region exhibiting length variation in fragile X syndrome’, Cell, 65(5), pp. 905–914.

Yasuma, T. et al. (2016) ‘Amelioration of diabetes by protein S’, Diabetes, 65(7), pp. 1940–1951.

Yokota, Y. et al. (2009) ‘The Adenomatous Polyposis Coli (APC) Protein is an Essential Regulator of Radial Glial Polarity and Construction of the Cerebral Cortex’, Neuron, 61(1), pp. 42–56.

Zalfa, F. et al. (2007) ‘A new function for the fragile X mental retardation protein in regulation of PSD-95 mRNA stability’, Nature Neuroscience, 10(5), pp. 578–587.

Zhang, M., Wang, Q. and Huang, Y. (2007) ‘Fragile X mental retardation protein FMRP and the RNA export factor NXF2 associate with and destabilize Nxf1 mRNA in neuronal cells’, Proceedings of the National Academy of Sciences, 104(24), pp. 10057–10062.

